# q2-longitudinal: a QIIME 2 plugin for longitudinal and paired-sample analyses of microbiome data

**DOI:** 10.1101/223974

**Authors:** Nicholas A. Bokulich, Yilong Zhang, Matthew Dillon, Jai Ram Rideout, Evan Bolyen, Huilin Li, Paul S. Albert, J. Gregory Caporaso

## Abstract

Studies of host-associated and environmental microbiomes often incorporate longitudinal sampling or paired samples in their experimental design. Longitudinal sampling provides valuable information about temporal trends and subject/population heterogeneity, offering advantages over cross-sectional and pre/post study designs. To support the needs of microbiome researchers performing longitudinal studies, we developed q2-longitudinal, a software plugin for the QIIME 2 microbiome analysis platform (https://qiime2.org). The q2-longitudinal plugin incorporates multiple methods for analysis of longitudinal and paired-sample data, including paired differences and distances, linear mixed effects models, microbial interdependence test, first differencing, and volatility analyses. The q2-longitudinal package (<u>https://github.com/qiime2/q2-longitudinal</u>) is open source software released under a BSD-3-Clause license and is freely available, including for commercial use.

## Introduction

Time is an important component in many microbiome studies. Sampling microbial communities repeatedly over time provides information on their development (1), their stability (2–4), or their response to and recovery from a treatment or disturbance (5–7). The frequency and scale of longitudinal sampling can range from pre-post studies, in which individual subjects are sampled before and after treatment, to long-term observation studies lasting months or years. Such studies benefit from the use of dynamic analytical methods, which evaluate trends over time in relation to one or more variables; paired methods, which evaluate the magnitude of change within individual subjects; and random effects models, which account for the variation inherent to complex biological systems (8). Cross-sectional studies that examine differences in the microbiome across environmental gradients (e.g., pH or temperature) can also benefit from many of the same techniques.

Longitudinal study of microbiomes presents several unique challenges. Many microbiome datasets (e.g., marker-gene and shotgun metagenome sequences) are compositional (relative abundance) data, which violate the assumptions of some conventional statistical methods (9). Incorporating phylogenetic distance information into longitudinal analyses can deepen insight into relationships between the microbiome and possible health and disease etiologies (10, 11). Specialized methods for microbiome analysis are becoming more prevalent as these issues become better understood (12, 13). However, many statistical and bioinformatics methods for longitudinal analyses of microbiomes can be difficult for non-specialists to implement and interpret (e.g., requiring programming skills or statistical knowledge), hindering wider adoption of appropriate methods.

To address the challenge of applying appropriate longitudinal testing in routine microbiome analyses, we developed q2-longitudinal (<u>https://github.com/qiime2/q2-longitudinal</u>), a suite of bioinformatics tools for paired and longitudinal microbiome analyses. This software package is a plugin for the microbiome bioinformatics platform QIIME 2 (<u>https://qiime2.org/</u>), and thus adopts the software architecture, multiple user interfaces (including a graphical user interface), provenance tracking, and other user benefits offered by QIIME 2. Many of the analyses provided in q2-longitudinal wrap pre-existing tools, streamlining their use and reducing the burden for users to install, run, and interpret. Other analyses adapt standard statistical approaches for microbiome data (non-parametric tests are used by default, but parametric equivalents are supported for some plugin actions). All analyses are provided as easy-to-use pipelines, generating publication-ready tables and plots that are generated by q2-longitudinal, adding additional value relative to using the underlying tools directly.

## Results and discussion

To demonstrate the methods currently available in q2-longitudinal, we present a re-analysis of data from the early childhood and the microbiome (ECAM) study (1). This study tracked the 16S rRNA gene microbiota composition of 43 infants in the U.S. sampled at regular intervals from birth to two years of age, and associations between antibiotic use, delivery mode, and predominant diet on microbiota composition and development.

### Paired differences between two time points

Longitudinal and paired-sample studies (e.g., pre-post studies) frequently test whether an experimental measurement, such as sample diversity or the abundance of a microbial species, differs between paired samples collected at two different time points. For continuous measurements, such as alpha diversity metrics (e.g., species richness within a sample), we can perform paired difference tests that compare the paired differences across subjects. Paired t-tests or Wilcoxon signed rank tests performed on these differences determine whether a population’s distribution changed in a consistent direction between the two time points. For comparing measurements or differences in measurements over time between groups, ANOVA or Kruskal-Wallis tests can be applied on the values or differences, respectively. These methods are implemented in q2-longitudinal’s “pairwise-differences” action.

As an example of “pairwise-differences”, we will test alpha diversity differences between sampling times in the ECAM study. During early childhood, gut bacterial species richness and other alpha diversity metrics increase dramatically from the basal levels observed at birth (1). We can use paired difference testing to demonstrate this effect, and to assess whether alpha diversity changes in subjects at different rates according to delivery mode. Paired differences in Shannon diversity at birth and 12 months of age are significant in subjects who are vaginally delivered (Wilcoxon false discovery rate-corrected p = 0.002) but not in cesarean-delivered subjects (p > 0.05) (Figure 1A). This is a potentially important finding: as alpha diversity generally increases dramatically over this period, the observation that cesarean-delivered subjects’ Shannon diversity does not increase may indicate that early disturbances (i.e., perinatal antibiotic use or bypassing microbial exposure in the birth canal) lead to stalled development of the microbiota in cesarean-delivered subjects (1).

**Figure 1.**
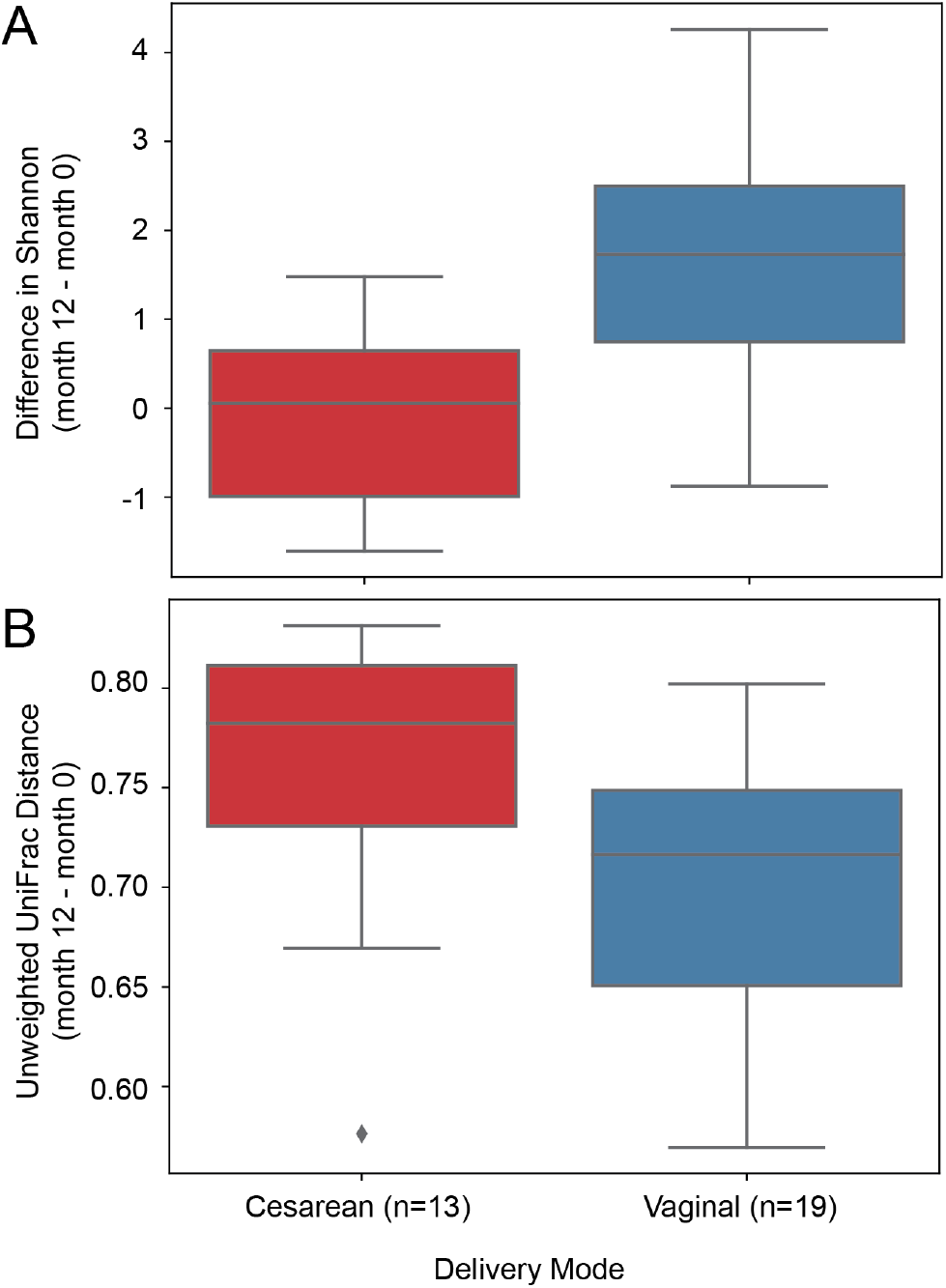
Paired difference (A) and paired distance tests (B) between birth and 12 months of age in vaginally delivered and cesarean delivered subjects. A, Bacterial Shannon diversity. B, Unweighted UniFrac distance. Boxplots show quartile distribution of differences or distances between each subject’s diversity at month 12 and month 0.

In addition to alpha diversity metrics, changes in sample ordination (e.g., on a principal coordinate plot), relative abundance (e.g., of a taxon, sequence variant, or operational taxonomic unit), or clinical data such as body mass index are other relevant candidates for this test. Raw difference data can also be easily exported in formats that are convenient for use in R, or other frameworks supporting more customized statistical tests.

### Paired distances between two time points

Longitudinal experiments and paired-sample experiments may also measure the degree of similarity or dissimilarity (beta diversity) between an individual’s microbiota community structure at two separate time points. This test is similar to the pairwise-differences described above, but in this case we examine the distances between sample pairs using a measure of community structure dissimilarity, such as UniFrac distance (14) or community distance metric. Statistical testing is performed with Wilcoxon rank sum tests or one-sample t-tests to determine whether one group’s microbiota differs across time (e.g., between pre- and post-treatment). This can be an important assessment of microbiota stability, discerning the impact of a disturbance or intervention on community structure or composition. This method is implemented in q2-longitudinal’s “pairwise-distances” action.

As an example of “pairwise-distances”, we will compute unweighted UniFrac distances between sampling times in the ECAM study. During the first few years of life, the microbiota changes rapidly before eventually stabilizing around 3 years of age (1). We can use paired distances to examine how this progression differs between infants delivered vaginally or by cesarean section. At 12 months of age, cesarean-born subjects’ microbiota compositions are more different from birth (i.e., unweighted UniFrac distances are greater) than vaginally born subjects’ microbiota (p = 0.010) (Figure 1B). This could indicate either greater instability in these subjects, or simply a larger change since birth as the microbiota of cesarean-delivered infants transitions to an age-typical community structure from a less typical starting point (1). Additional time points between birth and one year of age can help us to differentiate between these two possibilities with a more detailed longitudinal analysis.

### Analyzing longitudinal trajectories

Longitudinal experiments frequently measure observations over many time points, in which case paired differences or distances are inadequate to assess how the trajectory of a given metric changes over the course of an observation period. Investigators are often interested in measuring the impact of multiple factors on the trajectory of the dependent variable of interest; for example, the impact of a treatment and a subject’s sex on alpha diversity or species abundance measurements. When the longitudinal measurements are continuous, an approach commonly used to address such questions is linear mixed effects (LME) models (8). LME models examine the relationship between one or more independent variables (effects) and a single longitudinal response, where observations are made across dependent samples, e.g., in repeated-measures experiments. For example, a simple LME may include an intercept and slope term as both fixed and random effects. The fixed intercept and slope can be interpreted as the regression line for the average subject, while the random effects reflect individual departures from the average line for each subject. An attractive feature of the model is that it allows the investigator to explicitly model heterogeneity in the initial value and slope across subjects. This is important for longitudinal microbiome studies where we expect heterogeneity in the temporal pattern across individuals. Fixed effects are factors that may reflect group or time and assess the overall effect of the factor on the response. Random effects reflect variation in these effects across subjects. LME models are implemented in the “linear-mixed-effects” action in q2-longitudinal.

To demonstrate the use of the “linear-mixed-effects” action, let us revisit the question of alpha diversity changes during early life in the ECAM cohort. As samples were collected at roughly monthly intervals from these subjects, we can use LME to track the change in alpha diversity over time and in the context of multiple experimental variables; here we examine time and delivery as fixed effects, and apply a random intercept and slope as random effects (Figure 2, Table 1). Over the course of approximately 2 years, Shannon diversity significantly changes by age (months) (p < 0.001). Delivery mode and delivery mode X time interaction are both significant effects (p < 0.001). The statistically significant delivery mode effect provides evidence for a higher mean Shannon diversity at baseline for women undergoing Cesarean compared with vaginal delivery. However, the statistically significant mode X time interaction shows that the rate of change is higher for vaginal versus Cesarean delivery. Interestingly, these two features result in the crossing of the two trajectories demonstrating higher Shannon diversity after approximately 6 months of age. Delivery mode thus appears to be an important factor impacting gut microbiota diversity over the first two years of life, as previously described (1).

**Figure 2.**
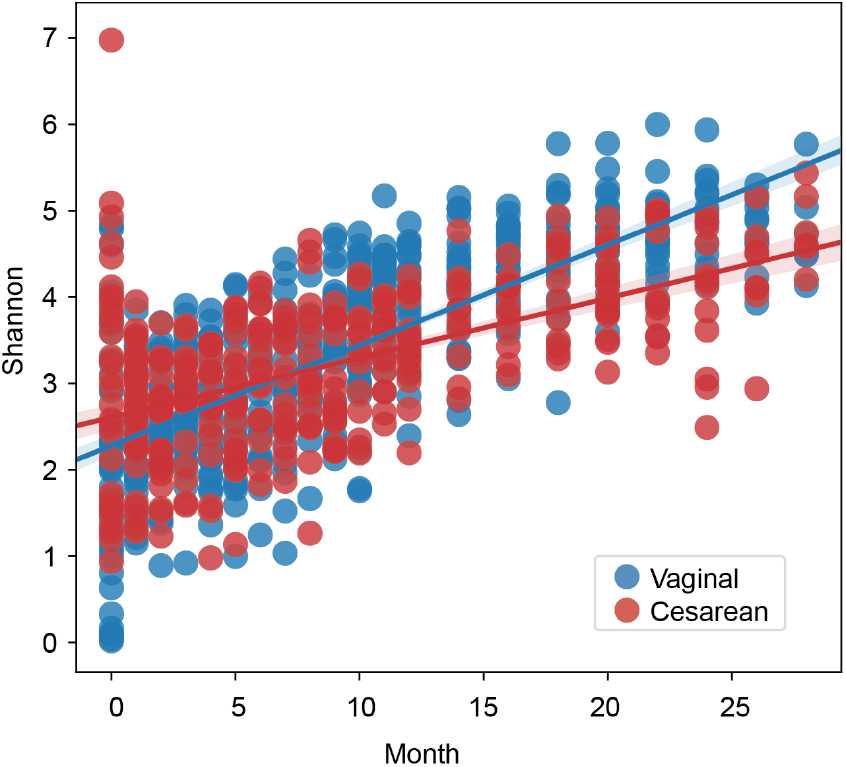
Scatterplots of Shannon diversity as a function of child age (months) and delivery mode. Linear regression trend lines are plotted for each group.

**Table 1.** Linear mixed effects model results for Shannon diversity in the ECAM study. Parameter estimates (coef.), standard error, Z score, and P value for each model parameter.

|  | Variables/Parameters | Estimates | Std.Err. | z | P Value |
| --- | --- | --- | --- | --- | --- |
| Fixed Effects | Intercept | 2.635 | 0.128 | 20.58 | < 0.001 |
|  | delivery[T.Vaginal] | -0.414 | 0.17 | -2.434 | 0.015 |
|  | month | 0.065 | 0.011 | 5.943 | < 0.001 |
|  | month:delivery[T.Vaginal] | 0.061 | 0.014 | 4.241 | < 0.001 |
| Random Effects | Intercept | 0.244 | 0.106 |  |  |
|  | Slope | -0.016 | 0.008 |  |  |
| Residual Error |  | 0.001 | 0.001 |  |  |

### Visualizing sample volatility

The temporal stability or volatility of a metric between individual subjects or groups of subjects can be an important measurement, indicating periods of disruption, disease, or abnormal events. Microbial volatility — the variance in microbial abundance, diversity, or other metrics over time — can be a marker of ecosystem disturbance or disease (4, 15, 16), and hence provides another important metric for comparison between experimental groups. We can visualize these fluctuations using control charts, which show how a variable changes over time in individuals or groups. These charts display “control limits” three standard deviations above and below the mean and “warning limits” two standard deviations above and below the mean to identify observations that deviate substantially from the mean. Observations at these time points could indicate aberrant conditions, e.g., due to disturbance or even sample contamination. Spaghetti plots, illustrating the longitudinal trajectory of each individual, support visual assessment of individual subjects’ stability, identifying aberrant individuals and time points.

Control plots with optional spaghetti plot overlays are supported in q2-longitudinal with the “volatility” visualizer (Figure 3A). Using these “volatility charts” to track Shannon diversity in the ECAM study, we see that vaginally and cesarean-delivered infants exhibit similar degrees of variance in Shannon diversity over time, but these groups exhibit divergent means immediately following birth and following one year of life (Figure 3A).

**Figure 3.**
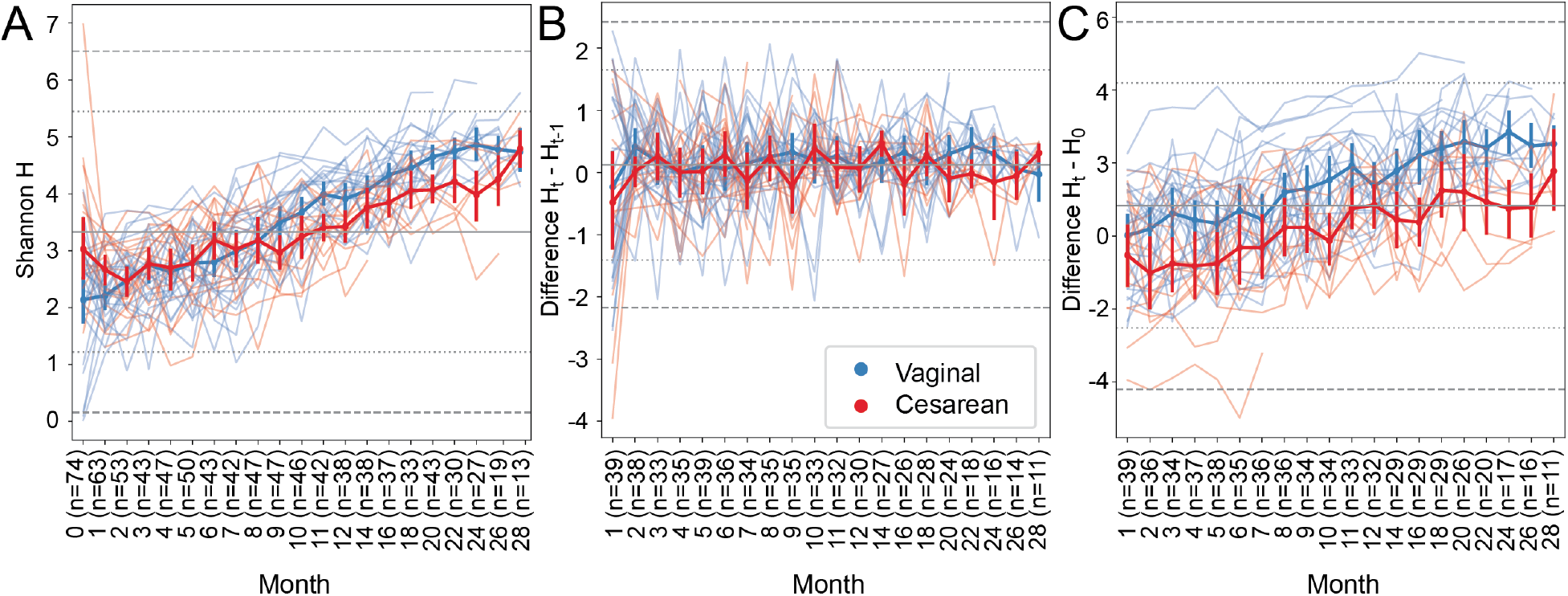
Volatility charts of Shannon H in ECAM subjects over time (A), rate of change (first differences) in Shannon H over time (B), and difference from baseline (C). Thick lines with error bars represent mean Shannon H (± standard deviation) for vaginally and cesarean-delivered subjects. Faded “spaghetti” lines represent the longitudinal trajectory for each individual subject. Horizontal lines represent the mean (solid midpoint) and two (dotted line) and three standard deviations from the mean (dashed line) computed across all samples. Sample sizes differ between subplots because some subjects in panel A were sampled more than once per month, and other subjects are missing samples for a particular month; replicates are dropped for first differencing, and missing samples affect the two adjacent time points. Note that x- and y-axis scales differ across the three plots to highlight difference in the scale that is most informative for each analysis.

### Computing first differences to track rate of change

Another way to view time series data is by assessing how the rate of change differs over time. We can do this through calculating first differences, which represent the magnitude of change between successive time points. If Yt is the value of metric Y at time t, the first difference at time t, ΔY_t_=Y_t_−Y_t−1_. This transformation is performed in the “first-differences” method in q2-longitudinal.

Figure 3B illustrates the first differences plot corresponding to longitudinal measurements of Shannon diversity shown in Figure 3A. Vaginally and cesarean-delivered infants show similar, high degrees of variance in rate of change in Shannon H, hovering around the mean ΔY = 0.121, indicating a very steady average rate of change over 28 months of life, with the exception of the first month of life when diversity slightly drops in both groups (Figure 3B). Such a constant difference is consistent with a linear relationship between time and Shannon H on the original scale (Figure 3A).

The “first-differences” method has an optional “baseline” parameter to instead calculate differences from a static point (e.g., baseline or a time point when a treatment is administered: ΔY_t_=Y_t_−Y_baseline_). Calculating baseline differences can help tease apart noisy longitudinal data to reveal underlying trends in individual subjects, such as some outlier subjects in the ECAM dataset (Figure 3C). In other cases, difference from baseline or treatment times could highlight significant experimental factors related to changes in diversity or other dependent variables.

A similar method implemented in q2-longitudinal is “first-distances”, which instead identifies the beta diversity (between-sample) distances between successive samples from the same subject based on a distance matrix. The output of first-distances is particularly empowering, because it allows us to analyze longitudinal changes in beta diversity using actions that cannot operate directly on a distance matrix, such as linear mixed effect models, or plotting with volatility charts (Figure 4). When applied to longitudinal changes in unweighted UniFrac distance (14) in the ECAM dataset, we see that vaginally and cesarean-delivered infants exhibit similar rates of phylogenetic transition (Figure 4A). This is marked by a dramatic shift in the first month of life, followed by gradual stabilization in the rate of change but a very large degree of variance. These groups only diverge in the first month of life (when cesarean-born infants exhibit a higher degree of change within individuals) and after two years of life (when sample sizes and statistical power are lower). Consistent with the close similarity between delivery modes, LME models indicate no significant differences between delivery modes, though month and intercept (p < 0.001) are both significant factors.

**Figure 4.**
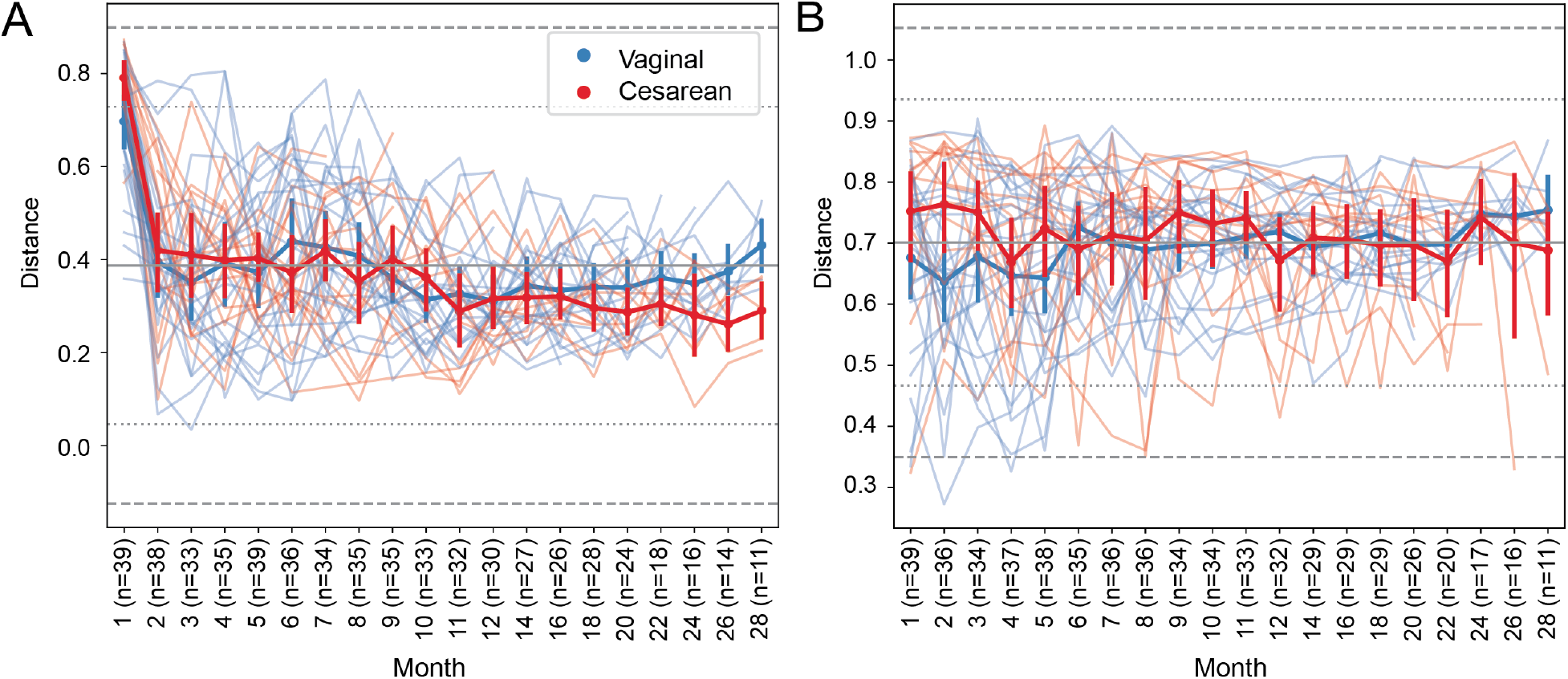
Volatility charts of longitudinal change in unweighted UniFrac distances between successive samples collected from the same subject in the ECAM dataset (A) and distance from baseline for each subject (B). Thick lines with error bars represent mean distance (± standard deviation) for vaginally and cesarean-delivered subjects. Faded “spaghetti” lines represent the longitudinal trajectory for each individual subject. Horizontal lines represent the mean (solid midpoint) and two (dotted line) and three standard deviations from the mean (dashed line) computed across all samples. Sample sizes differ between subplots because some subjects are missing samples for a particular month, resulting in fewer subjects eligible for first differencing at that month and the subsequent time point. Note that x- and y-axis scales differ across the three plots to highlight difference in the scale that is most informative for each analysis.

The “first-distances” method also has a “baseline” parameter for calculating distance from a static time point (Figure 4B), similar to the “first-differences” baseline parameter. This can be a useful approach for assessing how a subject differs from the start/end of a study, or from another static time point (e.g., to highlight fluctuations in community structure/composition related to a treatment) (Figure 4B).

### Quantifying shared features with first-distances

The first-distances method also allows us to track longitudinal change in the proportion of features that are shared between an individual’s samples. This can be performed by calculating pairwise Jaccard distance (proportion of features that are not shared) between each pair of samples with QIIME 2’s “diversity” plugin, and using first-distances to extract distances between successive samples (Figure 5A), or from baseline (Figure 5B). Applying this method to the ECAM dataset indicates that the intestinal microbiome becomes gradually more stable in the second year of life, as successive measurements exhibit less feature turnover and less variance in distance (Figure 5A). Distance from baseline remains high and volatile throughout the course of observation, indicating few shared taxa (Figure 5B).

**Figure 5.**
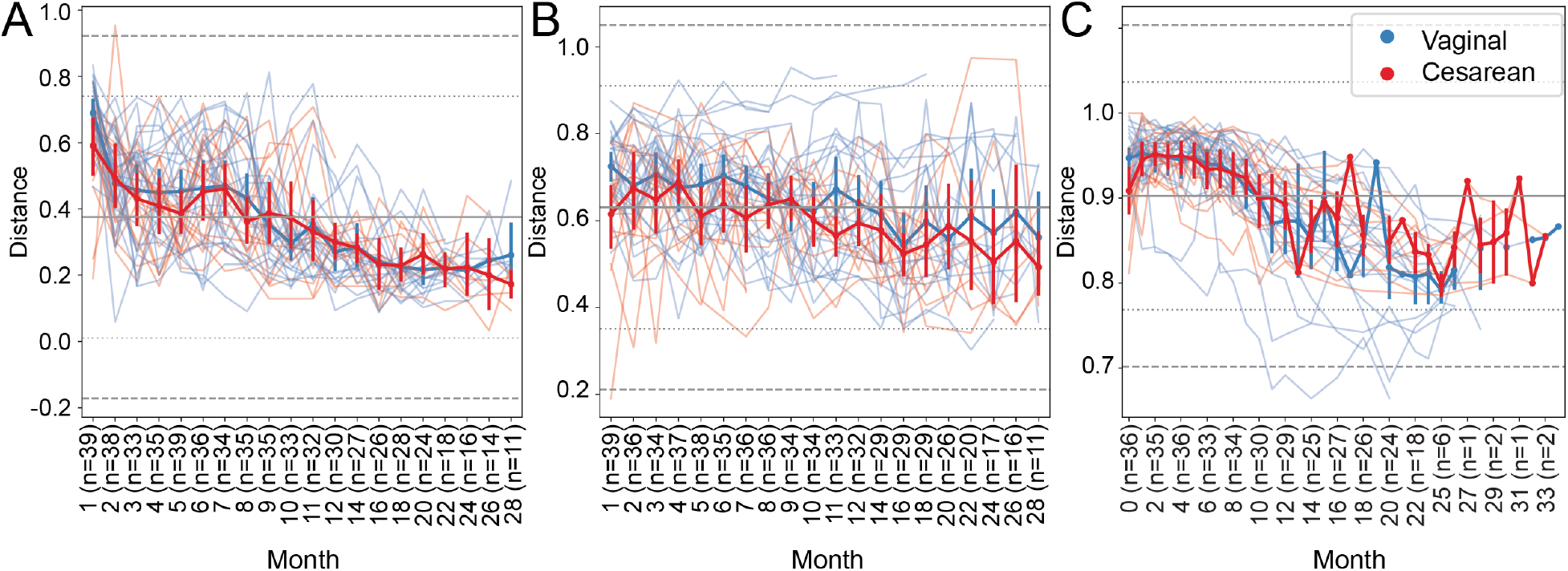
Volatility charts of longitudinal change in Jaccard distances between genus-level taxonomic compositions of successive samples collected from the same subject in the ECAM dataset (A), distance from baseline for each subject (B), and distance from their mother’s stool 16S rRNA sequence variant composition near birth (C). Thick lines with error bars represent mean distance (± standard deviation) for vaginally and cesarean-delivered subjects. Faded “spaghetti” lines represent the longitudinal trajectory for each individual subject. Horizontal lines represent the mean (solid midpoint) and two (dotted line) and three standard deviations from the mean (dashed line) computed across all samples. Sample sizes differ between subplots because some subjects are missing samples for a particular month, resulting in fewer subjects eligible for first differencing at that month and the subsequent time point. Note that x- and y-axis scales differ across the three plots to highlight difference in the scale that is most informative for each analysis.

The “baseline” parameter in first-distances and first-differences also provides the ability to track longitudinal change from a separate set of (non-longitudinal) samples. For example, we can track the number of shared features between the stool microbiota of infants and their mothers’ stool microbiota near the time of birth in the ECAM dataset (Figure 5C). Jaccard distance between sequence variant profiles indicates that very few variants are shared with a child’s mother during the first year of life, but distance decreases into the second year of life, when a higher proportion of sequence variants are shared between mother-infant dyads (Figure 5C). This indicates that as infants age they accumulate more microbiota characteristic of an adult gut ecosystem, as shown previously (1).

Using the “baseline” parameter to track longitudinal change from a static point opens many opportunities for comparisons common to some microbiome experimental designs. Other applications could include comparing similarity between the microbiota of human patients or gnotobiotic animals receiving fecal microbiota therapy and the composition of donor samples, between fermentations and their inocula, or between intact and disturbed environments during recovery from disturbance.

### Longitudinal analyses of feature data

Any action in q2-longitudinal that operates on sample metadata can also accept a feature table as input (i.e., abundance table of features per sample, e.g., sequence variants, taxa, genes, or metabolites), including “pairwise-differences”, “first-differences”, “volatility”, and “linear-mixed-effects”. To demonstrate these operations, we generated volatility charts and LME tests to track longitudinal changes in the relative abundance of genus *Bacteroides* in the ECAM study data. Volatility charts show that *Bacteroides* relative abundances are higher in vaginally delivered infants during the first year of life (Figure 6), as shown in the original study (1). A LME model of abundance vs. time and delivery mode with a random intercept and slope indicates a significant effect of both time and delivery mode on *Bacteroides* abundances (p < 0.001) (Table 2). The delivery mode X time interaction term is small and non-significant suggesting that the rate of change in relative abundance is constant across the two modalities.

**Figure 6.**
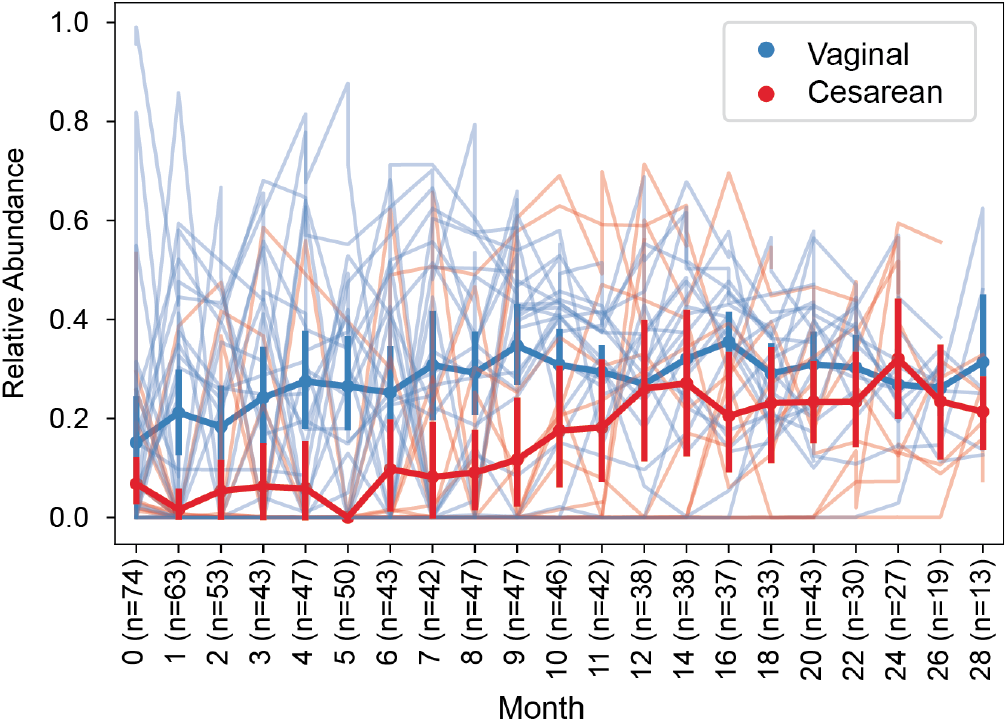
Spaghetti plot of longitudinal change in *Bacteroides* relative abundance in each subject in the ECAM dataset. Thick lines with error bars represent mean distance (± standard deviation) for vaginally and cesarean-delivered subjects. Faded “spaghetti” lines represent the longitudinal trajectory for each individual subject.

**Table 2.** Linear mixed effects model results for *Bacteroides* relative abundance in the ECAM study. Parameter estimates (coef.), standard error, Z score, and P value for each model parameter.

|  | Variables/Parameters | Estimates | Std.Err. | z | P Value |
| --- | --- | --- | --- | --- | --- |
| Fixed Effects | Intercept | 0.022 | 0.035 | 0.637 | 0.524 |
|  | delivery[T.Vaginal] | 0.184 | 0.046 | 3.972 | < 0.001 |
|  | month | 0.011 | 0.003 | 4.496 | < 0.001 |
|  | month:delivery[T.Vaginal] | -0.005 | 0.003 | -1.438 | 0.151 |
| Random Effects | Intercept | 0.019 | 0.033 |  |  |
|  | Slope | -0.001 | 0.002 |  |  |
| Residual Error |  | 0 | 0 |  |  |

### Temporal microbial interdependence

Within microbial communities, microbial populations do not exist in isolation but instead form complex ecological interaction webs. Whether these interdependence networks display the same temporal characteristics within subjects from the same group may indicate divergent temporal trajectories. To address this experimental question, q2-longitudinal implements the non-parametric microbial interdependence test (NMIT) (12). NMIT examines the longitudinal relationship between microbial features in a subject (e.g., taxa, sequence variants, or operational taxonomic units) to measure the similarity between individuals. The q2-longitudinal “nmit” method outputs a distance matrix containing the pairwise distances (microbial interdependence similarity) between individual subjects, which can then be assessed with other QIIME 2 methods such as PERMANOVA (17) tests to determine whether subject distances partition by phenotype or metadata categories, or principal coordinate analysis to visualize subject similarities.

Given the association of cesarean section with profound disturbances to the gut microbiome compared to vaginal delivery in the ECAM study (1), it can be expected that temporal microbial interdependence will be similarly impacted. Using NMIT to compute microbial interdependence and PERMANOVA (implemented in q2-diversity (https://github.com/qiime2/q2-diversity)) to test for an effect of delivery mode on between-subject distances, we find that delivery modes exhibit significantly different temporal microbial independence networks (p = 0.001) (Figure 7).

**Figure 7.**
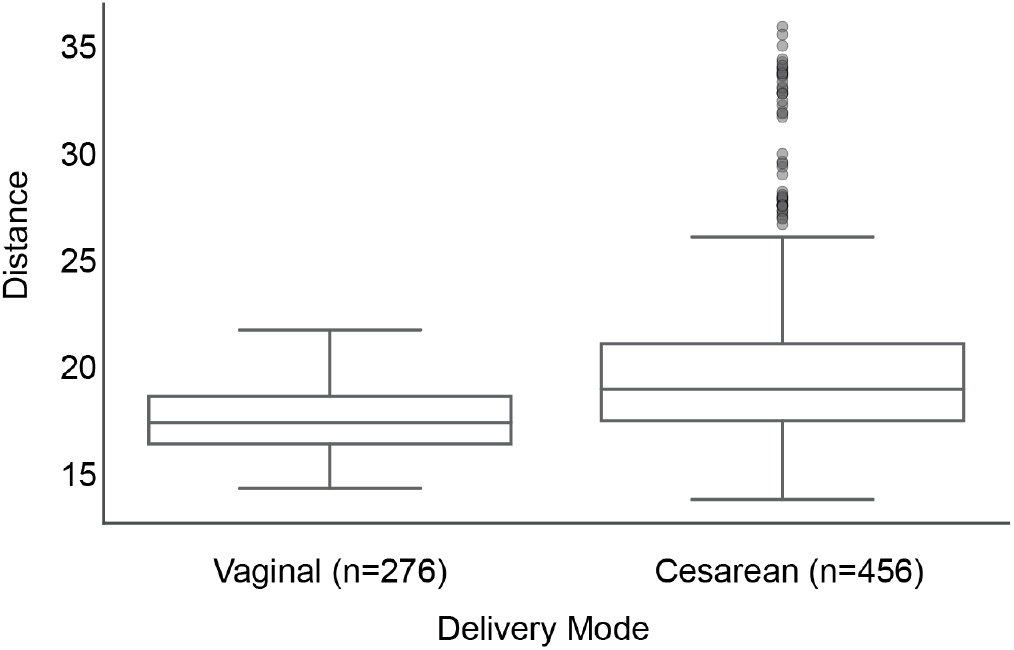
Temporal microbial interdependence distances between vaginally delivered subjects and other subjects in the ECAM study. Vaginally delivered subjects are more similar to each other than to cesarean-delivered subjects.

## Conclusions

Longitudinal designs for microbiome studies provide valuable information about time trends in biological activity. In addition, these designs allow investigators to distinguish between within-and between-subject variation, an important issue in characterizing heterogeneity in temporal patterns across experiments.

The q2-longitudinal package supports a variety of paired-sample and longitudinal tests relevant to studies of host-associated and environmental microbiomes. This includes methods for paired difference and distance testing, LME, microbial interdependence, analyses of volatility, and a variety of functions for generating publication-ready figures. Additional functions will be added to this plugin as they are developed (e.g., methods for tracking longitudinal volatility and shared species counts), and we welcome collaboration from other developers who would like their methods accessible through q2-longitudinal (get in touch on the QIIME 2 Forum at https://forum.qiime2.org/). This plugin is included in the QIIME 2 Core Distribution, and installation instructions and tutorials for the Core Distribution can be accessed at <u>https://qiime2.org</u>.

## Materials and methods

The q2-longitudinal package (<u>https://github.com/qiime2/q2-longitudinal</u>) is written in Python 3 and is accessible as a QIIME 2 plugin (<u>https://qiime2.org</u>). As a plugin in the QIIME 2 Core Distribution, users automatically have access to q2-longitudinal simply by installing QIIME 2, and can interact with the plugin using a variety of user interfaces (command line, Python API, and graphical user interfaces are included in the Core Distribution). The actions in this plugin utilize scipy (<u>https://scipy.org</u>), numpy (18), and pandas (19) for data manipulation and statistical testing, and matplotlib (20) and seaborn (<u>https://zenodo.org/record/12710</u>) for plotting. Tutorials and other information about the q2-longitudinal plugin are available at <u>https://qiime2.org</u>. This package is released under a BSD-3-Clause license and is freely available, including for commercial use.

### Test data

We use study data from the ECAM study (1) to demonstrate the features of q2-longitudinal. Raw sequence data (study id: 10249) were downloaded from Qiita (<u>http://qiita.microbio.me</u>) and analyzed with QIIME 2. Raw sequences were quality-filtered using DADA2 (21) to remove phiX, chimeric, and erroneous reads. Sequence variants were aligned using MAFFT (22) and used to construct a phylogenetic tree using FastTree 2 (23). Beta diversity was estimated using unweighted UniFrac distance (14). All other analyses were performed using q2-longidutinal.

Analysis data and notebooks used to generate all results in this study are available at <u>https://github.com/caporaso-lab/longitudinal-notebooks</u>.

## Acknowledgments

This project was funded in part by NSF Award 1565100, The Partnership for Native American Cancer Prevention (NIH/NCI U54CA143924 and U54CA143925), and an Arizona Board of Regents grant to JGC. The study was supported in part by NIH grants R01DK110014 to HL.

